# How do fluctuating temperatures alter the cost of development?

**DOI:** 10.1101/2023.05.19.541426

**Authors:** Amanda K Pettersen, Andreas Nord, Geoffrey While, Tobias Uller

**Affiliations:** The University of Sydney; Lund University; University of Tasmania

**Keywords:** developmental cost theory, hatching, incubation, metabolic rate, temperature, wall lizard, yolk

## Abstract

1. Quantifying how variable temperature regimes affect energy expenditure during development is crucial for understanding how future thermal regimes may impact early life survival and population persistence.
2. Developmental Cost Theory (DCT) suggests that there is an optimal temperature (T_opt_) that minimises energy expenditure during development (the “cost of development”). Exposure to fluctuating temperatures around an average of T_opt_ are anticipated to increase either development time or metabolic rate. As a result, embryos will rapidly deplete yolk reserves, and consequently hatch at a smaller size or with less residual yolk to support postnatal survival and growth.
3. Here, we studied total embryonic energy expenditure (development time and rate of CO_2_ production) and conversion of yolk into tissue in common wall lizards (*Podarcis muralis*) under three incubation treatments anticipated, based on DCT, to increase the cost of development: no variance (T_opt_ constant, 24 °C), low variance (22 – 26 °C), and high variance (18 – 30 °C).
4. As predicted, we found that increasing variance around T_opt_ increased the cost of development, despite reducing time to hatching. As a consequence, embryos on average hatched with 59% lower residual yolk reserves under high variance versus constant incubation temperatures.
5. Our results highlight how the relative temperature sensitivities of development time and metabolic rate determine the cost of development, which in turn may predict the ability of egg-laying ectotherms to persist in variable environments. We show that DCT can provide a mechanistic framework for understanding the widespread, but often seemingly idiosyncratic, effects of fluctuating incubation temperatures on hatchling tissue and residual yolk mass.

## Introduction

The development of an embryo from a single-celled zygote to a nutritionally independent hatchling is a critical life stage that is likely to be an important bottleneck for survival under increasingly extreme and variable environments. In egg-laying species, embryonic development is fuelled by maternally-provisioned yolk (Blaxter, 1969; Deeming & Ferguson, 1991; Mousseau & Fox, 1998). Energy derived from yolk is used for crucial processes during prenatal development including cell division and differentiation, growth, and maintenance, while residual yolk at hatching can be internalised to support postnatal growth and survival (Marshall et al., 2020; Murakami et al., 1992; Troyer, 1983; Vidal et al., 2002). The total energy expenditure during the period of embryonic development, also known as the “cost of development”, will be a product of 1) development time (time from fertilisation to hatching; *D*), and 2) rate of energy expenditure (metabolic rate; *MR*) (Pettersen et al., 2019). Any factor that increases either *D* or *MR* during prenatal development is anticipated to deplete yolk reserves which may in turn reduce residual yolk and/or the amount of tissue at hatching, with likely fitness consequences.

In ectotherms, temperature affects both developmental and metabolic rates (Pettersen et al., 2019). Developmental cost theory (DCT) posits that the thermal sensitivity of development time (*D_T_*) relative to the thermal sensitivity of metabolic rate (*MR_T_*) determines how the overall cost of development (*D* × *MR*) scales with temperature (Marshall et al., 2020). If the slope of *D_T_* and *MR_T_* functions differ, as shown for many ectotherm species, there will be an optimal temperature that minimises energy consumption during development (T_opt_), and thereby maximises residual yolk upon completion of development (Pettersen et al., 2019). At temperatures below T_opt_, *D_T_* > *MR_T_* and development time is extended to a greater extent than metabolic rate is reduced (Figure 1). Beyond T_opt_, *D_T_* < *MR_T_* and metabolic rate increases more than development time decreases with temperature. Using experimental data across seven phyla, DCT has been used to predict T_opt_ for 71 ectotherm species (Marshall et al., 2020). The effects of temperature on embryonic development is crucial for predicting the effect of changing temperature on population persistence, yet it remains poorly understood how well DCT predicts energy utilisation within species (Du et al., 2019).

**Figure 1.**
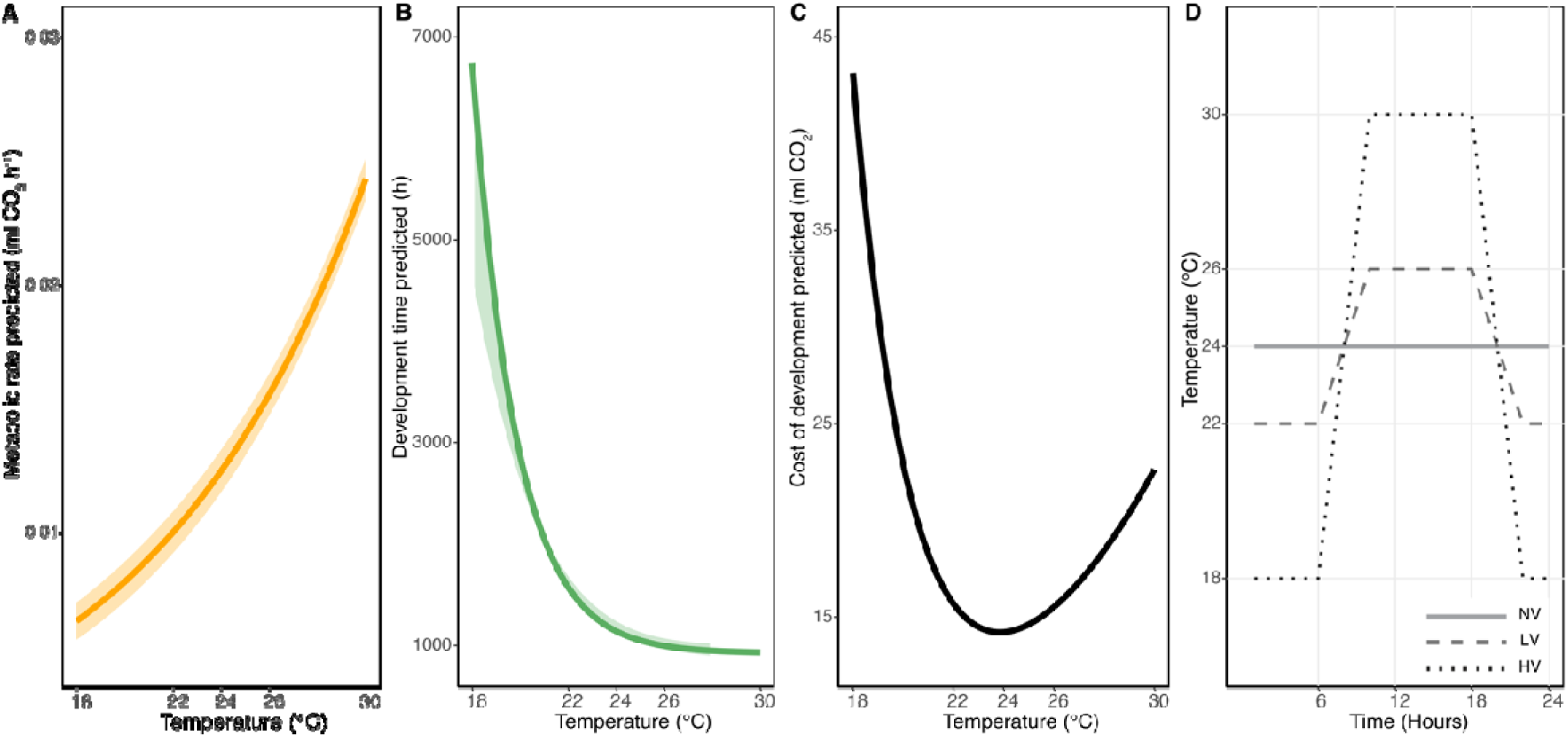
Graphical representation of *P. muralis* embryonic **A)** Temperature dependence of metabolic rate (ml CO_2_ h^-1^ ± 95% confidence interval; CI); *MR_T_* based on previous work (Pettersen et al., 2022), and **B)** Temperature dependence of development time (incubation time: number of hours from oviposition to hatching ± 95% CI); *D_T_* (Pettersen et al., 2022; While et al., 2015), used to calculate **C)** Cost of development; *C*_T_ (total CO_2_ production from oviposition until hatching). Grey vertical lines represent the temperature that minimises the cost of development (T_opt_, 24 °C) and the stable temperature used for the “no variance” (NV) treatment. Light grey dashed lines represent temperature range of “low variance” (LV: 22 – 26 °C) and black dotted lines represent temperature range of “high variance” (HV: 18 – 30 °C) treatments used in this study. Given that *C*_T_ is a convex function, embryos developing under variable temperatures are predicted to have greater cost of development than embryos developing under constant 24 °C. **D)** Programmed incubator settings over a 24h cycle throughout development for the three temperature treatments: NV (light grey line), LV (dashed grey line), and HV (dotted black line).

Natural environments are thermally variable, and it is well-known that the relationship between performance and temperature is often nonlinear in ectotherms (Bernhardt et al., 2018; Denny, 2017). While DCT predicts that incubation at temperatures that deviate from T_opt_ should increase the cost of development, the extent to which fluctuating temperatures will exacerbate total energy expenditure, due to an increase in *D*, *MR*, or both, has yet to be empirically tested. The extent to which embryos experiencing variable temperatures expend more energy to complete development compared to those developing under constant temperatures, will depend on the shape of the cost-temperature function, even when variable and constant temperatures have the same average temperature (Slein et al., 2023). For offspring, time spent developing at temperatures that deviate from T_opt_ therefore necessitates that they will either need to sacrifice energy reserves, hatching with lower residual yolk, or compromise the conversion of yolk into tissue, by hatching at a smaller size. Elucidating the effect of developmental temperatures that vary around a mean of T_opt_ on both the cost of development, and embryonic phenotypes, is thus crucial for understanding how well DCT predicts energy acquisition and allocation under fluctuating temperatures.

Egg-laying lizards provide a useful system to test DCT under fluctuating, ecologically relevant temperatures since egg incubation temperatures are variable in the field, often cycling through a consistent diel regime during incubation (Oufiero & Angilletta Jr., 2010a). Many lizard species hatch with a significant amount of residual yolk that is internalized before hatching – this energy resource can sustain offspring for several days post-hatching, and is likely to be under strong selection when food availability is low or unpredictable (Radder et al., 2004; Troyer, 1987; but see Radder et al. 2007). On the other hand, converting a greater proportion of yolk reserves towards tissue production may be beneficial if larger hatchling size confers greater survival, sprint speed, or reproductive output later in life (Sinervo, 1990; Sorci & Clobert, 1991; Uller & Olsson, 2010). Past studies have measured the effect of fluctuating versus constant incubation temperatures on hatchling phenotypes in lizards, including effects on embryonic development (see recent review by Raynal et al., (2022)), and metabolic rate (Du & Shine, 2010; Hall & Warner, 2020; Oufiero & Angilletta Jr., 2010b). Yet no studies have tested how deviations from the theoretical T_opt_ influence total energy expenditure and the conversion of yolk into tissue during development. This makes it difficult to assess patterns of developmental plasticity in relation to DCT, and their likely fitness implications.

Here, we estimated total energy expenditure (*D* × *MR*) and embryonic conversion of yolk into tissue in common wall lizards (*Podarcis muralis*) from three locally adapted altitude regions to test DCT under ecologically relevant, fluctuating incubation temperatures (Pettersen et al., 2023). First, we used existing measures of *D_T_* and *MR_T_* to estimate the constant temperature that minimises the cost of development (T_opt_) for *P. muralis*. Second, we measured how developmental cost varies under incubation temperatures that fluctuate around T_opt_ using three temperature regimes. All three temperature regimes had the same average temperature (24 °C; optimal for this species in terms of energy expenditure, Figure 1) but differed in the daily magnitude of fluctuation around this average (low: 24 ± 2 °C, high: 24 ± 6 °C). Third, we measured hatchling wet mass, dry tissue, and residual yolk mass across the three incubation temperatures regimes. Based on DCT we predicted that higher fluctuations around T_opt_ would impose a higher energetic cost of development, necessitating embryos that developed under fluctuating temperatures to either (*i*) compromise energy reserves at hatching and hatch at the same mass but with less residual yolk, or (*ii*) compromise conversion of yolk to tissue and hatch at a smaller size, than embryos developing at T_opt_.

## Materials and methods

### Calculation of P. muralis embryo thermal optima (T_opt)_

All analyses were conducted in R version 4.0.3 (R Core Team, 2023). We used development (incubation) time (*D*; Pettersen et al., <u>(2023; While et al., 2015</u>) and metabolic rate (*MR*; Pettersen et al., (2023)) data previously collected on Italian *P. muralis* populations, to calculate the temperature that minimises the cost of development (*C*) according to developmental cost theory (DCT), where:

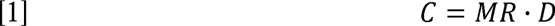

Exponential models were fit using the “nls” function to determine parameter estimates for *MR* and *D*, whereby:

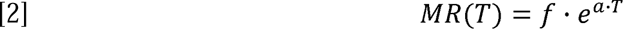

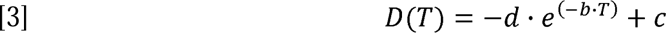

where *a* and *b* represent the temperature (*T*) dependencies of *MR* and *D* respectively, and *f*, *d*, and *c* are additional parameters that combined describe the decrease of development time, and increase of metabolic rate, with increases in incubation temperature. Based on the combined functions of [2] and [3], the temperature that minimises predicted *C* (referred to as “T_opt_”) was calculated to be 23.9 °C (Figure 1; see Supporting Information for details).

To estimate the expected increase in *C* under temperatures that fluctuate evenly around T_opt_ over a “low variance” (T_opt_ ± 2 °C) interval, we find the average value of function [8] (see Supporting Information) over 22 – 26 °C given by

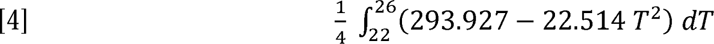

where development under low variance is predicted to cost 15.1 ml CO_2_, equivalent to a 4.1% increase in *C* from T_opt_. The expected increase in *C* when incubation temperature fluctuates evenly around T_opt_ over a “high variance” (T_opt_ ± 6 °C) interval, the average value of function [8] (see Supporting Information) over 18 – 30 °C is given by

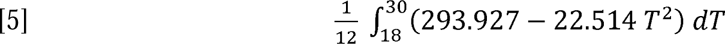

where development under high variance is predicted to cost 20.0 ml CO_2_, resulting in a 38% increase in *C* compared with development at T_opt_.

### Study system, field collection, and animal husbandry

The common wall lizard (*Podarcis muralis* Laurenti, 1768) is a small (50-70mm snout-to-vent length) oviparous lacertid. Adult, gravid females (n = 62) and adult males (n = 38) captured in the field during the prior reproductive season across three regions in central Italy: low-, mid-, and high-altitude (for details, see Pettersen et al., (2023)), were reared in animal facilities at Lund University, Sweden. During the reproductive season from April – June 2020, lizards were housed in a temperature-controlled room (24 °C day/20 °C night) with a light cycle of 12 L:12 D. One to two females and one male were housed in cages (590 × 390 × 415 mm) containing a sand substrate, bricks and logs as shelter, a sand box for egg laying, and a water bowl. Each cage contained a basking lamp (60 W) switched on for 8 h per day to create a thermal gradient (24 °C – 40 °C) and a UV light (EXOLTERRA 10.0 UVB fluorescent tube) for 6 h per day. Lizards were fed mealworms or crickets daily. Sand boxes were inspected daily for clutches to ensure egg collection within 24 h post-oviposition.

### Incubation treatments

Upon egg collection, we weighed the mother and eggs to the nearest 0.01g (Ohaus Scout SKX Precision Balance). Both females in the cage were visually inspected for evidence of having recently laid a clutch. Over the reproductive season, females laid between 1-5 clutches (mean: 3.48 ± SD: 1.09) with clutch sizes ranging 1-10 eggs (mean: 4.6 ± SD: 1.8). For the purpose of this experiment, we only sampled the 1^st^ and 2^nd^ clutches. We used a split clutch design to allocate eggs across three incubation treatments: one constant at T_opt_, “no variance” (NV; 24 °C), and two fluctuating thermal regimes with a mean temperature of 24 °C and either “low variance” (LV; ± 2°C, fluctuating between 22-26 °C) or “high variance” (HV; ± 6°C, fluctuating between 18-30 °C) to determine whether fluctuating temperatures that still maintain a mean T_opt_ will alter the energetic cost of development as manifested at the organismal level by smaller hatchling size, less residual yolk, or both. Fluctuating temperature regimes within this range will be encountered in the wild because of daily variation in soil temperatures (see also While et al. 2015 and Supporting Information in Pettersen et al., (2023)). Incubators (Snijders Labs Micro Clima-Series Economic Lux Chamber) were programmed on a 24h cycle: 9h at the low temperature of the treatment range (LV:22 °C/HV:18 °C), followed by a 3h ramping period to the upper treatment range (LV:26 °C/HV:30 °C) maintained for 9h and an equivalent ramping down, or a continuous 24 °C for NV treatment (Figure 1d). Eggs were incubated in small plastic containers filled two-thirds with moist vermiculite (5:1 vermiculite to water ratio) and sealed with clingfilm throughout development (except when they were temporarily removed for metabolic rate measurements, see details below) until hatching. The level at which inferences were made is individual developing embryos, however the level at which each treatment was applied is individual females (since some females contributed more than one egg to a given treatment). The number of females used per treatment is provided in Table 1, and the number of embryos measured for each treatment per female is provided in Table S1.

**Table 1.**
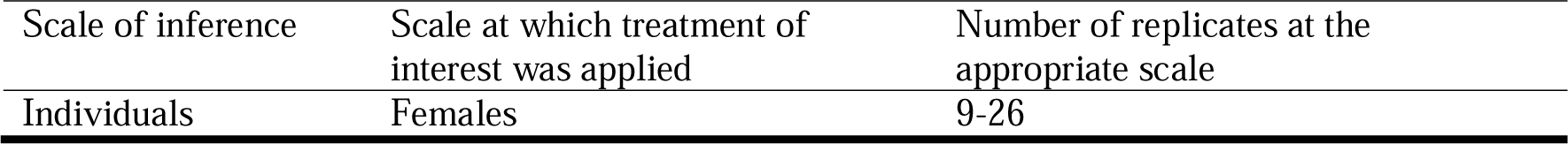
Replication Statement. for the scale of inferences made, the scale at which the treatment was applied, and the number of replicated for each treatment (exact numbers per measurement provided in Table S1).

### Measurement of embryonic metabolic rate

Pilot data from dissections on a subsample of first clutches was used to estimate days post oviposition (dpo) for embryos when they were 75% through development (NV: 38 ± 1 dpo, LV: 37 ± 2 dpo, HV: 35 ± 1 dpo). Embryo metabolic rate (*MR*) was then measured at 75% of development for the temperatures at which each embryo was exposed to within their treatment (NV:24 °C/LV: 22 °C, 24 °C, 26 °C/HV:18 °C, 24 °C, 30 °C; Figure 1a). For LV and HV, measures were taken in increasing order of temperatures, reflecting the daily ramping of incubation temperature. Embryos were kept in measurement vials with the lid removed for 60 min prior to sealing for metabolic rate measurements, which ensured they were in thermal equilibrium with the measurement temperature when the experiment started. Hence, metabolic rates were unlikely to depend on the direction of the ramping profile (as shown previously in Nord and Folkow). To determine whether *MR* remained broadly consistent across development, we also measured NV embryos at 25% (11-14 dpo) and 50% (23-26 dpo) through development (at 24 °C only). Rate of CO_2_ production, or V CO_2_, was calculated from measures of fractional CO_2_ using closed-system, stop-flow respirometry as per methods described previously (Cordero et al., 2017; Pettersen et al., 2022). In brief, individual eggs were placed into 60ml glass chambers containing 50ml moist vermiculite. Each chamber was then connected to and flushed with dry scrubbed pressurised air (using drierite desiccant; Hammond Drierite at a flow rate of 200 (±20) ml min^-1^ (standard temperature and pressure, dry; STPD; registered using a FM-8 mass flow meter from Sable Systems International, Las Vegas, NV, USA). The sealed vials were then placed into incubators set at the measurement temperature. After 180 min, a baseline measurement of CO_2_ was taken before re-connecting the chambers to the airflow. Excurrent air (at approximately 200 ml min^-1^, STPD) from the chambers was dried using magnesium perchlorate (Mg(ClO_4_)_2_), after which we measured the total volume of CO_2_ accumulated in the chambers using a FC-10 CO_2_ analyser (Sable Systems International). Every embryo MR was measured at each temperature within their respective treatment. V CO_2_ in ml h^-1^ was calculated using the *metabR* package (Noble, 2021), by integrating the area under the %CO_2_ curve above the baseline.

### Development time, cost of development, and hatchling trait measurements

Eggs were checked daily for signs of hatching to determine development time; the number of hours from oviposition until hatching (*D*). Since hatching success was high across all treatments (NV: 100%, LV: 99.4%, HV: 98.7%) we did not analyse survival. The cost of development (total ml CO_2_ produced) by each individual was calculated as V CO_2_ at each temperature (measured within each treatment) multiplied by the time in hours spent at each temperature. Within 24 h of hatching, individuals were humanely euthanised and measures of hatchling wet mass (to 0.01g), hatchling dry mass and yolk dry mass (to 0.01 mg), snout-to-vent length (svl; mm), and total length (mm) were taken. All hatchling condition measurements were carried out by the same author. Since hatchling wet mass includes residual yolk, it is not an informative metric for quantifying total energy allocation towards tissue synthesis. The ability to dissect residual yolk from the hatchling allows for direct measurement of the allocation of initial yolk towards tissue production and residual yolk, under temperature regimes that are more or less costly in terms of energy expenditure. Hence, hatchling tissue and yolk were placed into a drying oven at 60 °C for 3 days, then weighed using a microbalance (Sartorius Research 200D). Further drying did not lead to any further reductions in sample weights (AKP personal observation).

### Hypothesis tests

To quantify how variation in incubation temperature around T_opt_ (24 °C) alters the overall energy cost of development, and conversion of yolk to tissue during development, we fitted linear mixed effects models for the following response variables: (1) metabolic rate (*MR*, measured as V CO_2_), (2) development time (*D*, measured as time from oviposition until hatching), (3) cost of development (*C*, as a function of temperature-dependent *MR* x total time spent at each temperature until hatching), (4) wet mass at hatching, (5) dry mass at hatching and (6) dry residual yolk mass at hatching. All models were fit using the “lmer” function within the *lme4* package (Bates et al., 2015). Fixed predictor variables of incubation ‘Treatment’ (NV, LV, HV), ‘Egg mass’, ‘Region’ of parent origin (high, mid, low altitude), as well as two-way interactions among these, and ‘Clutch order’ (1-2; for *D* and wet mass at hatching), were tested. For *MR*, we first tested for differences among treatments at the intermediate temperature (24 °C). We then ran separate models for each treatment since *MR* was measured at different temperatures within each treatment, and ‘Temperature’ (and its interaction with Region) were tested. Maternal identity was included as a random effect in all models. To account for non-independent replication when eggs from the same female were included within a treatment, we used the average value per clutch (Harrison et al., 2018). Egg mass and metabolic rate were natural log transformed for all hypothesis tests, to satisfy the assumption of heteroscedasticity, and to allow for meaningful interpretation of scaling patterns (Glazier, 2021; Niklas & Hammond, 2014).

### Model selection and parameter estimation

Candidate models (including predictor variables Treatment, Log_n_ Egg mass, Region, and their interactions, summarised in Table 2) were ranked using Akaike Information Criterion (AIC) averaged for those with Δ conditional Akaike information criterion (AICc) <4 using the R package *MuMin* v.1.43.17 (Barton, 2009). Post-hoc tests using the *emmeans* (Lenth, 2022) package were run to compare pairwise differences in fixed factors retained in the final models.

**Table 2.** Summary of candidate models used in model selection for 1. Log_n_ Metabolic rate, 2. Log_n_ Development time, 3. Log_n_ Cost of Development, 4. Hatchling wet mass, 5. Hatchling tissue dry mass, and 6. Hatchling residual yolk dry mass. Fixed predictor variables were Temperature (within each Treatment: LV = 22, 24, 26 °C, HV = 18, 24, 30 °C), Log_n_ Egg mass, Region (High, Mid, Low) and Clutch order (1^st^, 2^nd^). Mother ID was included as a random effect for all models. Clutch order is included in models for development time and hatchling wet mass (only 2^nd^ clutches were measured for MR, C, and hatchling dry mass and residual yolk mass). Note for Response 1 (Log_n_ Metabolic rate), Temperature was not included for models of NV treatment measured at a single temperature (24 °C).

| Response | Model no. | Parameters |
| --- | --- | --- |
| 1 | 1 | Temperature + 1 |
| 1 | 2 | Temperature + Log<sub>n</sub> Egg mass |
| 1 | 3 | Temperature + Log<sub>n</sub> Egg mass + Region |
| 1 | 4 | Temperature + Log<sub>n</sub> Egg mass + Region + Temperature x Log<sub>n</sub> Egg mass |
| 1 | 5 | Temperature + Log<sub>n</sub> Egg mass + Region + Temperature x Region |
| 1 | 6 | Temperature + Log<sub>n</sub> Egg mass + Region + |
|  |  | Log<sub>n</sub> Egg mass x Region |
| 2,3,4,5,6 | 7 | Treatment + 1 |
| 2,3,4,5,6 | 8 | Treatment + Log<sub>n</sub> Egg mass |
| 2,3,4,5,6 | 9 | Treatment + Log<sub>n</sub> Egg mass + Region |
| 2,4 | 10 | Treatment + Log<sub>n</sub> Egg mass + Region + Clutch order |
| 2,4 | 11 | Treatment + Log<sub>n</sub> Egg mass + Region + Clutch order +<br>Treatment x Log<sub>n</sub> Egg mass |
| 2,4 | 12 | Treatment + Log<sub>n</sub> Egg mass + Region + Clutch order +<br>Treatment x Region |
| 2,4 | 13 | Treatment + Log<sub>n</sub> Egg mass + Region + Clutch order +<br>Log<sub>n</sub> Egg mass x Region |
| 3,5,6 | 14 | Treatment + Log<sub>n</sub> Egg mass + Region +<br>Treatment x Log<sub>n</sub> Egg mass |
| 3,5,6 | 15 | Treatment + Log<sub>n</sub> Egg mass + Region +<br>Treatment x Region |
| 3,5,6 | 16 | Treatment + Log<sub>n</sub> Egg mass + Region +<br>Log<sub>n</sub> Egg mass x Region |

## Results

### (i) Metabolic rate

Metabolic rate (*MR*; CO_2_) measured at 24 °C (T_opt_) was significantly higher in the high variance (HV; 18-30 °C, mean ± SD: 1.51·10^-2^ ± 2.75 ·10^-3^ ml CO_2_ h^-1^) and low variance (LV; 22-26 °C, mean ± SD: 1.38·10^-2^ ± 2.76 ·10^-3^ ml CO_2_ h^-1^) treatments compared with the no variance treatment (NV; 24 °C, mean ± SD: 1.26·10^-2^ ± 2.00 ·10^-3^ ml CO_2_ h^-1^; *F*_2,168_ = 23.49, *p* <0.0001). Within LV and HV treatments, *MR* increased with measurement temperature (Table 3, Figure 2). We found a significant effect of log_n_ Egg mass on log_n_ *MR* for the NV treatment, but this relationship was non-significant for LV and HV (Table 3). *MR* measured at 24 °C increased approximately 1.6x across development, but this was only significantly different between 25% (mean ± SD: 8.10·10^-3^ ± 2.29 ·10^-3^ ml CO_2_ h^-1^) and 75% (mean ± SD: 1.26·10^-2^ ± 2.00 ·10^-3^ ml CO_2_ h^-1^) completion of development. There was no observed interaction between development stage and egg mass, indicating consistent metabolic scaling across early ontogeny (Table S3, Figure S1).

**Figure 2.**
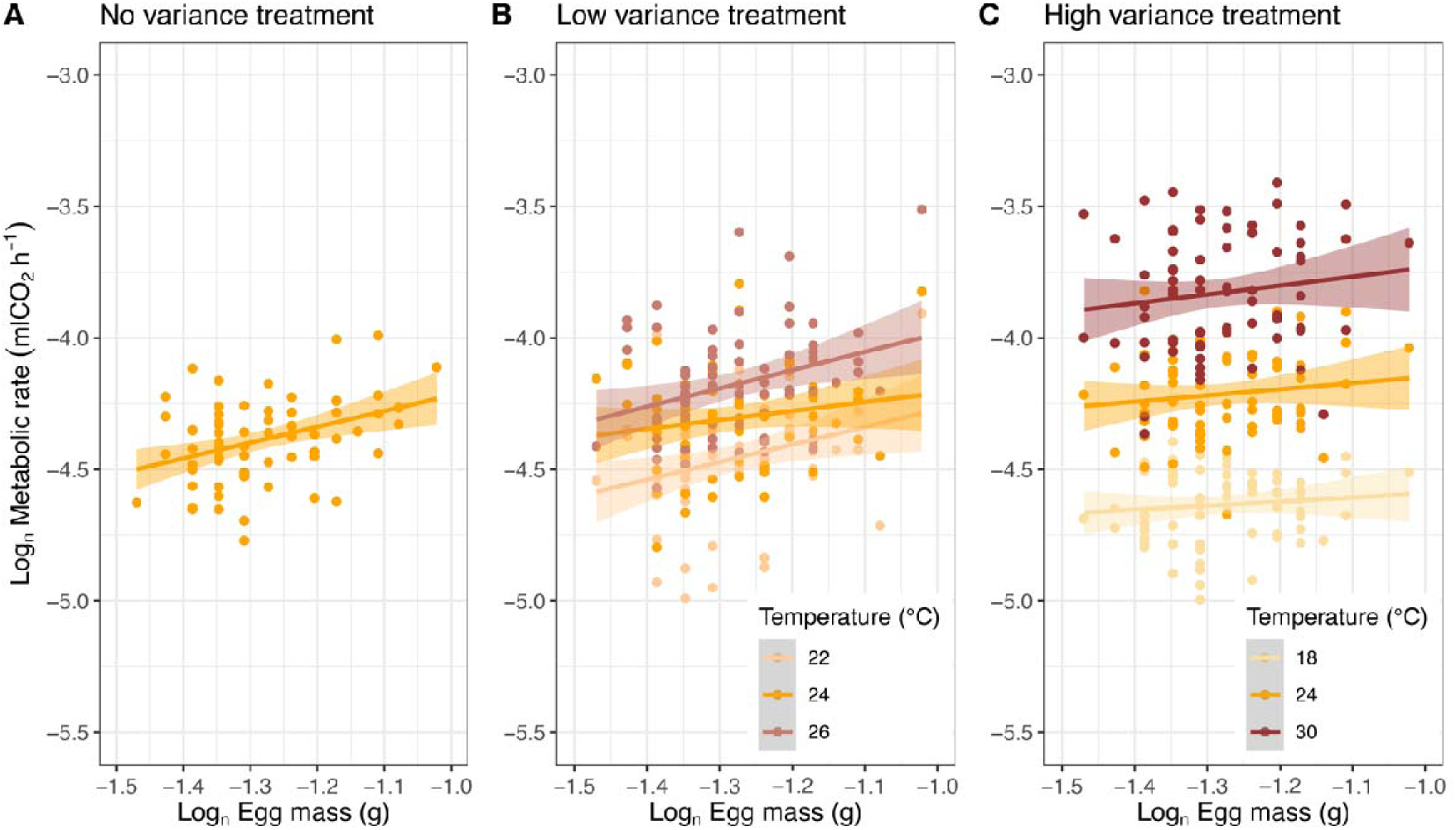
Response of Metabolic rate (Log_n_ CO_2_; ml CO_2_ h^-1^), to temperature across three incubation treatments: **A)** constant, “no variance” (NV; measured at 24 °C), **B)** low variance (LV; measured at 22 °C, 24 °C, 26 °C), and **C)** high variance (HV; measured at 18 °C, 24 °C, 30 °C). Data points show raw data and lines are fitted to individual responses with a single line fitted using a linear model. Orange data points are measures of metabolic rate at T_opt_ (24 °C). Lighter and darker colours represent deviations below and above T_opt_, respectively.

**Table 3.** Summary of model output (fixed and random effects) and significance tests and marginal means of fixed effects from best-fitting linear mixed effects models describing the relationship between 1. Metabolic rate, 2. Development time, 3. Cost of development, 4. Wet mass, 5. Dry mass, 6. Dry residual yolk mass, with predictor variables: Temperature (within treatment), Treatment (NV = no variance, LV = low variance, HV = high variance), Log_n_ Egg mass, Region, and Clutch order, and their interactions as shown in Table 2. Final model parameters were included based on model selection (AICc) shown in Table S2. Significant estimates shown in bold: \**p* < 0.05, \*\**p* < 0.01, \*\*\**p* < 0.001. Letters represent significant pairwise differences across levels (a > b > c). Note that responses for 1, 2, and 3 are back transformed from model output log_n_ estimates.

| | Estimate | $\chi^2$ | df | $p$ | emmean (±SE) |
| --- | --- | --- | --- | --- | --- |
| **1a. Log<sub>n</sub> Metabolic rate (NV)** |  |  |  |  |  |
| Intercept | -3.58 | 200.80 | 1 | **<0.001\*\*\*** |  |
| Log<sub>n</sub> Egg mass | 0.63 | 10.21 | 1 | **<0.01\*\*** |  |
| Mother ID (random) | 0.09 | 0.92 | 1 | 0.34 |  |
| **1b. Log<sub>n</sub> Metabolic rate (LV)** |  |  |  |  |  |
| Intercept | -3.84 | 132.30 | 1 | **<0.001\*\*\*** |  |
| Temperature | 24 °C: 0.15<br>26 °C: 0.28 | 216.54 | 2 | **<0.001\*\*\*** | 22 °C: $1.19 \cdot 10^{-2}$ ($3.41 \cdot 10^{-4}$)<sup>c</sup><br>24 °C: $1.38 \cdot 10^{-2}$ ($3.96 \cdot 10^{-4}$)<sup>b</sup><br>26 °C: $1.58 \cdot 10^{-2}$ ($4.51 \cdot 10^{-4}$)<sup>a</sup> |
| Log<sub>n</sub> Egg mass | 0.46 | 3.15 | 1 | 0.08 |  |
| Mother ID (random) | 0.18 | 132.66 | 1 | **<0.001\*\*\*** |  |
| **1c. Log<sub>n</sub> Metabolic rate (HV)** |  |  |  |  |  |
| Intercept | -4.60 | 256.09 | 1 | **<0.001\*\*\*** |  |
| Temperature | 24 °C: 0.42<br>30 °C: 0.80 | 1530.51 | 2 | **<0.001\*\*\*** | 18 °C: $9.75 \cdot 10^{-3}$ ($1.99 \cdot 10^{-4}$)<sup>c</sup><br>24 °C: $1.47 \cdot 10^{-2}$ ($3.71 \cdot 10^{-4}$)<sup>b</sup><br>30 °C: $2.17 \cdot 10^{-2}$ ($5.49 \cdot 10^{-4}$)<sup>a</sup> |
| Log<sub>n</sub> Egg mass | 0.03 | 0.01 | 1 | 0.91 |  |
| Mother ID (random) | 0.14 | 82.09 | 1 | **<0.001\*\*\*** |  |
| **2. Log<sub>n</sub> Development time** |  |  |  |  |  |
| Intercept | 7.04 | 2003063.25 | 1 | **<0.001\*\*\*** |  |
| Treatment | NV: 0.10<br>LV: 0.10 | 711.43 | 2 | <0.001*** | NV: 1257.65 (1.00)<sup>a</sup><br>LV: 1251.38 (1.00)<sup>a</sup><br>HV: 1132.29 (1.00)<sup>b</sup> |
| Mother ID (random) | 0.03 | 127.98 | 1 | <0.001*** |  |
| **3. Log<sub>n</sub> Cost of development** |  |  |  |  |  |
| Intercept | 3.43 | 274.05 | 1 | <0.001*** |  |
| Treatment | NV: -0.12<br>LV: -0.04 | 26.16 | 2 | <0.001*** | NV: 15.64 (1.02)<sup>b</sup><br>LV: 16.95 (1.02)<sup>a</sup><br>HV: 17.64 (1.02)<sup>a</sup> |
| Log<sub>n</sub> Egg mass | 0.44 | 7.40 | 1 | <0.01** |  |
| Mother ID (random) | 0.09 | 23.15 | 1 | <0.001*** |  |
| **4. Hatchling wet mass** |  |  |  |  |  |
| Intercept | 0.79 | 816.09 | 1 | <0.001*** |  |
| Treatment | NV: 0.02<br>LV: 0.02 | 45.03 | 2 | <0.001*** | NV: 3.49·10<sup>-1</sup> (3.93·10<sup>-3</sup>)<sup>a</sup><br>LV: 3.43·10<sup>-1</sup> (3.92·10<sup>-3</sup>)<sup>a</sup><br>HV: 3.33·10<sup>-1</sup> (3.93·10<sup>-3</sup>)<sup>b</sup> |
| Log<sub>n</sub> Egg mass | 0.36 | 273.23 | 1 | <0.001*** |  |
| Mother ID (random) | 0.02 | 42.92 | 1 | <0.001*** |  |
| **5. Hatchling tissue dry mass** |  |  |  |  |  |
| Intercept | 0.16 | 281.35 | 1 | <0.001*** |  |
| Treatment | NV: 1.40·10<sup>-7</sup><br>LV: 4.05·10<sup>-4</sup> | 3.54 | 2 | 0.17 | NV: 6.76·10<sup>-2</sup> (9.21·10<sup>-4</sup>)<sup>a</sup><br>LV: 6.88·10<sup>-2</sup> (9.16·10<sup>-4</sup>)<sup>a</sup><br>HV: 6.74·10<sup>-2</sup> (9.21·10<sup>-4</sup>)<sup>a</sup> |
| Log<sub>n</sub> Egg mass | 0.08 | 99.46 | 1 | <0.001*** |  |
| Mother ID (random) | 5.32·10<sup>-3</sup> | 46.26 | 1 | <0.001*** |  |
| **6. Hatchling residual yolk dry mass** |  |  |  |  |  |
| Intercept | 1.03·10<sup>-3</sup> | 62.65 | 1 | <0.001*** |  |
| Treatment | NV: 3.76·10<sup>-4</sup><br>LV: 1.84·10<sup>-4</sup> | 9.35 | 2 | <0.01** | NV: 1.41·10<sup>-3</sup> (1.30·10<sup>-4</sup>)<sup>a</sup><br>LV: 1.22·10<sup>-3</sup> (1.29·10<sup>-4</sup>)<sup>ab</sup><br>HV: 1.03·10<sup>-2</sup> (1.30·10<sup>-4</sup>)<sup>b</sup> |
| Mother ID (random) | 6.82·10<sup>-4</sup> | 51.13 | 1 | <0.001*** |  |

### (ii) Development time

Embryos incubated at HV (18-30 °C) hatched, on average, five days earlier (HV mean ± SD: 1142.54 ± 54.23 hours) than embryos incubated in LV (HV mean ± SD: 1250.17 ± 46.94 hours) and NV (HV mean ± SD: 1262.48 ± 46.94 hours) treatments, while no significant differences in development time between LV and NV were observed (Table 3, Figure 3a). We observed no significant effect of egg mass, region, clutch order (or interactions of these) on development time (Table S2).

**Figure 3.**
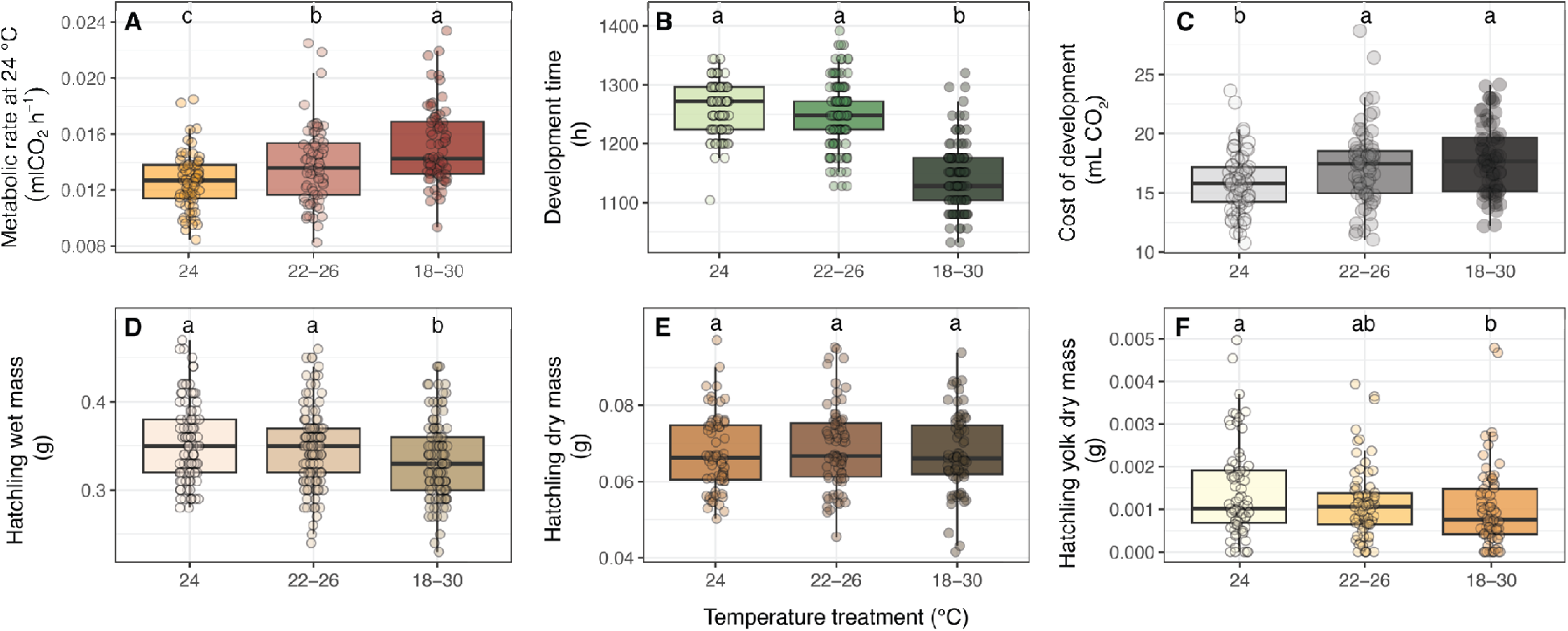
Tukey’s boxplots for response variables: a) Metabolic rate (measured at 24 °C), b) Development time, c) Cost of development, d) Hatchling wet mass (tissue and yolk mass combined), e) Hatchling tissue dry mass, and f) Hatchling residual yolk dry mass, to incubation temperature treatment (constant, “no variance” (NV; 24 °C; light shading), low variance (LV; 22-26 °C) and high variance (HV; 18-30 °C; dark shading)). Data points show raw data. Letters represent significant pairwise differences across levels (a > b > c) based on model output (note: raw data shown does not account for embryo mass included in all models).

### (iii) Cost of development

The total CO_2_ production to complete development, calculated from the temperature-specific V CO_2_ (*MR*) at 75% of development and total development time (*D*) increased with incubation temperature variation (Table 3). Based on the final best-fitting model, the ‘cost of development’ for an average size egg (0.28 g) was lowest under NV treatment (mean ± SD: 15.87 ± 2.54 ml CO_2_), intermediate for LV (mean ± SD: 17.19 ± 3.24 ml CO_2_), and highest for HV (mean ± SD: 17.84 ± 2.96 ml CO_2_). This reflected an increase in cost of 8% and 12% above NV for LV and HV, respectively. however no significant differences between LV and HV were found (*p* = 0.07, Figure 3b). The cost of development did not differ across regions (Tables S2 and S3).

### (iv) Wet mass at hatching

Hatchling wet mass (tissue and residual yolk; g) was significantly higher in the NV (mean ± SD: 0.36 ± 0.05g) and LV (mean ± SD: 0.35 ± 0.04g) treatments, compared with HV (mean ± SD: 0.34 ± 0.05g; Table 3, Figure 3c). Wet mass at hatching was significantly positively correlated with initial egg mass, however there was no significant difference in hatchling wet mass between regions or clutch order (Table S2). For an average size egg (0.28g) we calculated that embryos incubated at NV hatched 6% and 2% heavier than those under HV and LV, respectively.

### (v) Dry hatchling and yolk mass

Analysis of the dry mass of hatchlings and residual yolk revealed that hatchling dry mass was consistent across the incubation temperature treatments (Table 3 and S3, Figure 3d). We found that on average, embryos incubated at constant 24 °C (NV; mean ± SD: 1.41·10^-3^ ± 1.09·10^-3^g) hatched with 20% and 59% more yolk reserves, compared with the LV (mean ± SD: 1.14·10^-3^ ± 8.38·10^-4^g) and HV (mean ± SD: 1.02·10^-3^ ± 9.56·10^-4^g) respectively. However, there were no significant difference in dry yolk mass between LV and HV (Table 3, Figure 3e). While larger eggs resulted in hatchlings with higher tissue dry mass, there was no effect of egg mass on residual yolk dry mass at hatching (Table S2).

## Discussion

Many regions of the globe are experiencing altered patterns of temperature variability (Buckley & Huey, 2016; Wang & Dillon, 2014). Here, we show evidence that thermal variability may exacerbate the cost of development for egg-laying ectotherms. Our results demonstrate that, in wall lizards, increasing variation around the optimal incubation temperature (T_opt_) increases the cost of development such that hatchlings utilise more residual yolk and consequently begin life outside the egg with fewer energetic reserves. The extent to which these effects of fluctuating temperatures around T_opt_ will impact fitness is likely to be context dependent. Despite exacerbating developmental cost, there may still be selection for shorter development times that potentially increase metabolic rate, however this relationship was not observed in our study. If faster development does incur greater overall energy cost, then it may only be selected for when resources in the postnatal environment are abundant such that any fitness consequences of high costs of development are offset. However, where the postnatal environment is characterised by low or unpredictable resource availability then the consequences of the costs of development may be severe. Indeed, environments that have greater fluctuating thermal and hydric regimes on a seasonal basis, often show equally strong seasonality in food availability (Gilbert & Miles, 2016). Hence, thermally variable environments may present a twofold cost for developing embryos: a greater cost of development and therefore greater immediate reliance on external sources of food as well as a reduced availability of those external resources upon hatching.

As predicted, embryo energy expenditure (cost of development) was higher under fluctuating versus stable incubation treatments. This expectation was based on the convex shape of the temperature-cost function – which is a product of both metabolic rate (*MR*) and development time (*D*). Based on data from constant incubation temperatures, we anticipated that embryos exposed to thermal variance would show higher developmental energy costs, relative to embryos incubated under constant conditions, potentially due to extended development time, increased metabolic rates, or both. While we observed that exposure to high fluctuating temperatures increased overall metabolic rates, it also caused embryos to hatch sooner than those under constant or low fluctuating temperatures. This may help to explain why our predicted increase in cost of development above NV (4% and 38% for LV and HV respectively) differed from the actual increase (8% and 12%). For *P. muralis* experiencing thermal variability during development, exposure to temperatures above and below T_opt_ *decreased* development time to a lesser extent than it *increased* overall metabolic rates, resulting in overall increased energy expenditure with increasing time spent at temperatures further away from the optimum.

Due to the nonlinear relationship between development time and temperature, we expected a positive net effect of fluctuating temperatures on development time yet observed a decrease in *D* under fluctuating temperatures. Faster development under fluctuating temperatures has also been observed in other reptile species (Du & Shine, 2010; Fischer et al., 2011; Georges et al., 2005; Hall & Warner, 2020; Lin et al., 2008; Lu et al., 2009; Shine, 2004; Shine & Harlow, 1996) and may be due to physiological plasticity (acclimatisation), whereby embryos respond to shifts in temperature and compensate *D* accordingly. Alternatively, faster development under fluctuating temperatures may be a result of the thermal inertia of embryo body temperature cooling after exposure to the upper environmental temperature, which may also explain why higher metabolic rates are maintained in the thermal variance treatments when embryos were measured at 24 °C. Investigating the complicated responses of thermal history on embryo physiology, and how this may alter predicted implications of environmental change, is an important avenue of future research (Denny, 2019; Du & Shine, 2010; Pottier et al., 2022).

While DCT predicts that fluctuating temperatures will commonly impose energetic costs to developing embryos, what has been less clear, is *how* this increased cost will be paid for. In *P. muralis*, the increased cost of development under fluctuating temperatures did not affect the conversion of yolk into tissue but did result in lower energy reserves at hatching. This suggests that wall lizard embryos may prioritise tissue production over hatchling energy reserves when costs of development are high. There is evidence that internalised residual yolk reserves at hatching act as an insurance mechanism that can sustain hatchlings in the event of low foraging success (Radder et al., 2004; Troyer, 1983, 1987). On average, embryos incubated under high (± 6 °C) and low (± 2 °C) temperature variability around T_opt_ (24 °C) hatched with 59% and 20% lower yolk reserves, respectively, compared to incubation at constant T_opt_. This suggests that hatching at sufficient size, regardless of the energy needed to attain it, may be of greater importance for post-hatching fitness components such as survival and growth than retaining greater energy reserves. The importance of preserving yolk relative to synthesising tissue is likely to be context dependent, yet measures of the consequences of this trade-off for survival in the field are lacking.

Incubation temperature produces strong and pervasive effects on post-hatching survival in ectotherms, that have been well documented in the literature, particularly in lizards (Noble, Stenhouse, Riley, et al., 2018; While et al., 2018). Mechanistic explanations have been put forward to explain these effects, yet few studies have considered the fitness consequences of energetic cost of development (Cordero et al., 2017; Ji & Braña, 1999; Oufiero & Angilletta Jr., 2010a), and measures of both the temperature-dependence of embryonic metabolic rates and developmental time are rare. DCT provides a framework for predicting how the mean and variation of incubation temperature will affect the cost of development, and through this, fitness. Based on the shape of the cost function (as a product of non-invasive measures of *MR_T_* and *D_T_*), deviations in incubation temperature from T_opt_, including fluctuations around a mean of T_opt_, will increase developmental cost. However, exactly how much an increase in a degree fluctuation of incubation temperature will cost in terms of energy expenditure depends on the average temperature (as illustrated in Figure 1). This observation may help to explain the highly variable responses of *D* and *MR* under fluctuating incubation temperatures that have been reported in ectotherms (Bozinovic et al., 2011; Niehaus et al., 2012; Slein et al., 2023).

While we recognise the diversity of temperature effects on embryogenesis, we propose that many of the gross effects on size, mass, and energy consumption that have been observed in ectotherms (Noble, Stenhouse, & Schwanz, 2018; While et al., 2018) will be well explained by the temperature-dependence of metabolic rate relative to development time. Combining these measures, while also collecting data on the consequences of these costs in the wild, represents an important step towards understanding the mechanisms underlying the effects of developmental temperature on survival.

Both metabolic and developmental rates can vary between populations, reflecting either local adaptation (Conover & Schultz, 1995; Demarco, 1992; Pettersen, 2020) or plasticity (Cordero et al., 2017; Gangloff et al., 2019; Macagno et al., 2018). In wall lizards, developmental rate shows counter-gradient variation, whereby embryos from cooler, high altitude and latitude populations develop faster than those from warm climates when placed under common garden conditions (Pettersen et al., 2022; While et al., 2015). In our study, we found that population differences had a marginal influence on the optimal temperature from an energy-efficiency perspective – embryos from different climatic regimes were found to generate similar estimates of the cost of development and did not differ in their residual yolk at hatching. Under the assumption that the cost of development may vary across a lineage (Marshall et al. 2020), we can predict that populations that occupy the hot or cold extremes of the lineage’s distribution will therefore pay a higher cost. This raises the possibility that DCT may be able to predict species ranges based on mean and variation in nest temperatures. Complementary to relying on estimates of embryo thermal tolerance to inform species distributions (e.g., Kearney and Porter (2009)), it is plausible that measures of fitness quantified through the cost of development, can provide accurate predictions required to assess the impact of increasing thermal variance on the response of populations under global change.

## References

Barton, K. (2009). MuMIn: Multi-model inference. *Http://R-Forge.r-Project.Org/Projects/Mumin/*. https://cir.nii.ac.jp/crid/1572824499154168192

Bates, D., Mächler, M., Bolker, B., & Walker, S. (2015). Fitting Linear Mixed-Effects Models Using lme4. Journal of Statistical Software, 67(1), Article 1. 10.18637/jss.v067.i01

Bernhardt, J. R., Sunday, J. M., Thompson, P. L., & O’Connor, M. I. (2018). Nonlinear averaging of thermal experience predicts population growth rates in a thermally variable environment. Proceedings of the Royal Society B: Biological Sciences, 285(1886), 20181076. 10.1098/rspb.2018.1076

Blaxter, J. H. S. (1969). 4 Development: Eggs and Larvae. In Fish Physiology (Vol. 3, pp. 177–252). Elsevier. 10.1016/S1546-5098(08)60114-4

Bozinovic, F., Bastías, D. A., Boher, F., Clavijo-Baquet, S., Estay, S. A., & Angilletta, M. J. (2011). The Mean and Variance of Environmental Temperature Interact to Determine Physiological Tolerance and Fitness. Physiological and Biochemical Zoology, 84(6), 543–552. 10.1086/662551

Buckley, L. B., & Huey, R. B. (2016). Temperature extremes: Geographic patterns, recent changes, and implications for organismal vulnerabilities. Global Change Biology, 22(12), 3829–3842. 10.1111/gcb.13313

Conover, D. O., & Schultz, E. T. (1995). Phenotypic similarity and the evolutionary significance of countergradient variation. Trends in Ecology & Evolution, 10(6), Article 6. 10.1016/S0169-5347(00)89081-3

Cordero, G. A., Andersson, B. A., Souchet, J., Micheli, G., Noble, D. W. A., Gangloff, E. J., Uller, T., & Aubret, F. (2017). Physiological plasticity in lizard embryos exposed to high-altitude hypoxia. Journal of Experimental Zoology Part A: Ecological and Integrative Physiology, 327(7), Article 7. 10.1002/jez.2115

Deeming, D. C., & Ferguson, M. W. J. (Eds.). (1991). *Egg Incubation: Its* Effects on Embryonic Development in Birds and Reptiles. Cambridge University Press. 10.1017/CBO9780511585739

Demarco, V. (1992). Embryonic Development Times and Egg Retention in Four Species of Sceloporine Lizards. Functional Ecology, 6(4), 436–444. 10.2307/2389281

Denny, M. (2017). The fallacy of the average: On the ubiquity, utility and continuing novelty of Jensen’s inequality. Journal of Experimental Biology, 220(2), Article 2. 10.1242/jeb.140368

Denny, M. (2019). Performance in a variable world: Using Jensen’s inequality to scale up from individuals to populations. Conservation Physiology, 7(1), coz053. 10.1093/conphys/coz053

Du, W.-G., & Shine, R. (2010). Why do the eggs of lizards (Bassiana duperreyi: Scincidae) hatch sooner if incubated at fluctuating rather than constant temperatures? Biological Journal of the Linnean Society, 101(3), 642–650. 10.1111/j.1095-8312.2010.01525.x

Du, W.-G., Shine, R., Ma, L., & Sun, B.-J. (2019). Adaptive responses of the embryos of birds and reptiles to spatial and temporal variations in nest temperatures. Proceedings of the Royal Society B: Biological Sciences, 286(1915), Article 1915. 10.1098/rspb.2019.2078

Fischer, K., Kölzow, N., Höltje, H., & Karl, I. (2011). Assay conditions in laboratory experiments: Is the use of constant rather than fluctuating temperatures justified when investigating temperature-induced plasticity? Oecologia, 166(1), 23–33. 10.1007/s00442-011-1917-0

Gangloff, E. J., Sorlin, M., Cordero, G. A., Souchet, J., & Aubret, F. (2019). Lizards at the Peak: Physiological Plasticity Does Not Maintain Performance in Lizards Transplanted to High Altitude. Physiological and Biochemical Zoology, 92(2), 189–200. 10.1086/701793

Georges, A., Beggs, K., Young, J. E., & Doody, J. S. (2005). Modelling Development of Reptile Embryos under Fluctuating Temperature Regimes. Physiological and Biochemical Zoology, 78(1), 18–30. 10.1086/425200

Gilbert, A. L., & Miles, D. B. (2016). Food, temperature and endurance: Effects of food deprivation on the thermal sensitivity of physiological performance. Functional Ecology, 30(11), Article 11. 10.1111/1365-2435.12658

Glazier, D. S. (2021). Biological scaling analyses are more than statistical line fitting. Journal of Experimental Biology, 224(11). 10.1242/jeb.241059

Hall, J. M., & Warner, D. A. (2020). Ecologically relevant thermal fluctuations enhance offspring fitness: Biological and methodological implications for studies of thermal developmental plasticity. Journal of Experimental Biology, 223(19), jeb231902. 10.1242/jeb.231902

Harrison, X. A., Donaldson, L., Correa-Cano, M. E., Evans, J., Fisher, D. N., Goodwin, C. E. D., Robinson, B. S., Hodgson, D. J., & Inger, R. (2018). A brief introduction to mixed effects modelling and multi-model inference in ecology. PeerJ, 6, e4794. 10.7717/peerj.4794

Ji, X., & Braña, F. (1999). The influence of thermal and hydric environments on embryonic use of energy and nutrients, and hatchling traits, in the wall lizards (Podarcis muralis). Comparative Biochemistry and Physiology Part A: Molecular & Integrative Physiology, 124(2), Article 2. 10.1016/S1095-6433(99)00111-7

Kearney, M., & Porter, W. (2009). Mechanistic niche modelling: Combining physiological and spatial data to predict species’ ranges. Ecology Letters, 12(4), 334–350. 10.1111/j.1461-0248.2008.01277.x

Lenth, R. V. (2022). Estimated Marginal Means, aka Least-Squares Means (R package version 1.7.4-1) [Computer software].

Lin, L.-H., Li, H., An, H., & Ji, X. (2008). Do temperature fluctuations during incubation always play an important role in shaping the phenotype of hatchling reptiles? Journal of Thermal Biology, 33(3), 193–199. 10.1016/j.jtherbio.2007.12.003

Lu, H.-L., Hu, R.-B., & Ji, X. (2009). The variance of incubation temperatures does not affect the phenotype of hatchlings in a colubrid snake, Xenochrophis piscator. Journal of Thermal Biology, 34(3), Article 3. 10.1016/j.jtherbio.2008.12.002

Macagno, A. L. M., Zattara, E. E., Ezeakudo, O., Moczek, A. P., & Ledón-Rettig, C. C. (2018). Adaptive maternal behavioral plasticity and developmental programming mitigate the transgenerational effects of temperature in dung beetles. Oikos, 127(9), Article 9. 10.1111/oik.05215

Marshall, D. J., Pettersen, A. K., Bode, M., & White, C. R. (2020). Developmental cost theory predicts thermal environment and vulnerability to global warming. Nature Ecology & Evolution, 4(3), Article 3. 10.1038/s41559-020-1114-9

Mousseau, T. A., & Fox, C. W. (1998). Maternal Effects As Adaptations. Oxford University Press.

Murakami, H., Akiba, Y., & Horiguchi, M. (1992). Growth and utilization of nutrients in newly-hatched chick with or without removal of residual yolk. Growth, Development, and Aging: GDA, 56(2), 75–84.

Niehaus, A. C., Angilletta, M. J., Jr, Sears, M. W., Franklin, C. E., & Wilson, R. S. (2012). Predicting the physiological performance of ectotherms in fluctuating thermal environments. Journal of Experimental Biology, 215(4), 694–701. 10.1242/jeb.058032

Niklas, K. J., & Hammond, S. T. (2014). Assessing Scaling Relationships: Uses, Abuses, and Alternatives. International Journal of Plant Sciences, 175(7), 754–763. 10.1086/677238

Noble, D. W. A. (2021). *MetabR/MR.R at master · daniel1noble/metabR*. GitHub. https://github.com/daniel1noble/metabR

Noble, D. W. A., Stenhouse, V., Riley, J. L., Warner, D. A., While, G. M., Du, W.-G., Uller, T., & Schwanz, L. E. (2018). A comprehensive database of thermal developmental plasticity in reptiles. Scientific Data, 5(1), Article 1. 10.1038/sdata.2018.138

Noble, D. W. A., Stenhouse, V., & Schwanz, L. E. (2018). Developmental temperatures and phenotypic plasticity in reptiles: A systematic review and meta-analysis. Biological Reviews, 93(1), Article 1. 10.1111/brv.12333

Nord, A., & Folkow, L. P. (2018). Seasonal variation in the thermal responses to changing environmental temperature in the world’s northernmost land bird. Journal of Experimental Biology, 221(1), jeb171124. 10.1242/jeb.171124

Oufiero, C. E., & Angilletta Jr., M. J. (2010a). Energetics of Lizard Embryos at Fluctuating Temperatures. Physiological and Biochemical Zoology, 83(5), Article 5. 10.1086/656217

Oufiero, C. E., & Angilletta Jr., M. J. (2010b). Energetics of Lizard Embryos at Fluctuating Temperatures. Physiological and Biochemical Zoology, 83(5), 869–876. 10.1086/656217

Pettersen, A. K. (2020). A review and synthesis of countergradient thermal sensitivity of developmental rates in reptiles. Frontiers in Physiology.

Pettersen, A. K., Ruuskanen, S., Nord, A., Nilsson, J. F., Miñano, M. R., Fitzpatrick, L. J., While, G. M., & Uller, T. (2022). Population divergence in maternal investment and embryo energy use and allocation reveals adaptive responses to cool climates (p. 2022.12.07.519527). bioRxiv. 10.1101/2022.12.07.519527

Pettersen, A. K., Ruuskanen, S., Nord, A., Nilsson, J. F., Miñano, M. R., Fitzpatrick, L. J., While, G. M., & Uller, T. (2023). Population divergence in maternal investment and embryo energy use and allocation suggests adaptive responses to cool climates. Journal of Animal Ecology, n/a(n/a). 10.1111/1365-2656.13971

Pettersen, A. K., White, C. R., Bryson-Richardson, R. J., & Marshall, D. J. (2019). Linking life-history theory and metabolic theory explains the offspring size-temperature relationship. Ecology Letters, 22(3), Article 3. 10.1111/ele.13213

Pottier, P., Burke, S., Zhang, R. Y., Noble, D. W. A., Schwanz, L. E., Drobniak, S. M., & Nakagawa, S. (2022). Developmental plasticity in thermal tolerance: Ontogenetic variation, persistence, and future directions. Ecology Letters, 25(10), 2245–2268. 10.1111/ele.14083

R Core Team. (2023). R: A language and environment for statistical computing. R Foundation for Statistical Computing, Vienna, Austria. [Computer software]. https://www.R-project.org/.

Radder, R. S., Shanbhag, B. A., & Saidapur, S. K. (2004). Yolk partitioning in embryos of the lizard, Calotes versicolor: Maximize body size or save energy for later use? Journal of Experimental Zoology Part A: Comparative Experimental Biology, 301A(9), 783–785. 10.1002/jez.a.95

Raynal, R. S., Noble, D. W. A., Riley, J. L., Senior, A. M., Warner, D. A., While, G. M., & Schwanz, L. E. (2022). Impact of fluctuating developmental temperatures on phenotypic traits in reptiles: A meta-analysis. Journal of Experimental Biology, 225(Suppl_1), jeb243369. 10.1242/jeb.243369

Shine, R. (2004). Seasonal shifts in nest temperature can modify the phenotypes of hatchling lizards, regardless of overall mean incubation temperature. Functional Ecology, 18(1), 43–49. 10.1046/j.0269-8463.2004.00806.x

Shine, R., & Harlow, P. S. (1996). Maternal Manipulation of Offspring Phenotypes via Nest-Site Selection in an Oviparous Lizard. Ecology, 77(6), Article 6. 10.2307/2265785

Sinervo, B. (1990). The Evolution of Maternal Investment in Lizards: An Experimental and Comparative Analysis of Egg Size and Its Effects on Offspring Performance. Evolution, 44(2), 279–294. 10.1111/j.1558-5646.1990.tb05198.x

Slein, M. A., Bernhardt, J. R., O’Connor, M. I., & Fey, S. B. (2023). Effects of thermal fluctuations on biological processes: A meta-analysis of experiments manipulating thermal variability. Proceedings of the Royal Society B: Biological Sciences, 290(1992), 20222225. 10.1098/rspb.2022.2225

Sorci, G., & Clobert, J. (1991). Natural selection on hatchling body size and mass in two environments in the common lizard (Lacerta vivipara). Evolutionary Ecology Research, 1, 303–316.

Troyer, K. (1983). Posthatching yolk energy in a lizard: Utilization pattern and interclutch variation. Oecologia, 58(3), 340–344. 10.1007/BF00385233

Troyer, K. (1987). Posthatching Yolk in a Lizard: Internalization and Contribution to Growth. Journal of Herpetology, 21(2), 102–106. 10.2307/1564470

Uller, T., & Olsson, M. (2010). Offspring size and timing of hatching determine survival and reproductive output in a lizard. Oecologia, 162(3), Article 3. 10.1007/s00442-009-1503-x

Vidal, E. A. G., DiMarco, F. P., Wormuth, J. H., & Lee, P. G. (2002). Influence of temperature and food availability on survival, growth and yolk utilization in hatchling squid. Bulletin of Marine Science, 71(2), 915–931.

Wang, G., & Dillon, M. E. (2014). Recent geographic convergence in diurnal and annual temperature cycling flattens global thermal profiles. Nature Climate Change, 4(11), Article 11. 10.1038/nclimate2378

While, G. M., Noble, D. W. A., Uller, T., Warner, D. A., Riley, J. L., Du, W.-G., & Schwanz, L. E. (2018). Patterns of developmental plasticity in response to incubation temperature in reptiles. Journal of Experimental Zoology Part A: Ecological and Integrative Physiology, 329(4–5), 162–176. 10.1002/jez.2181

While, G. M., Williamson, J., Prescott, G., Horvathova, T., Fresnillo, B., Beeton, N. J., Halliwell, B., Michaelides, S., & Uller, T. (2015). Adaptive responses to cool climate promotes persistence of a non-native lizard. Proceedings of the Royal Society B-Biological Sciences, 282(1803), Article 1803. 10.1098/rspb.2014.2638

