## Supporting information for "How do fluctuating temperatures alter the cost of development?"

**Materials and methods**

*Calculation of P. muralis embryo thermal optima (T_opt_)*

Using previously collected data on the thermal sensitivity of *MR* and *D* (Pettersen et al., 2022; While et al., 2015), we estimated parameters for equations [2] and [3], where the thermal sensitivity of *MR* was found to be

[6]	$MR(T) =0.031{\cdot e}^{0.104\cdot T}$

while the relationship between development time and temperature was best fit by

[7]	$D(T)=2.674\cdot{10}^{6}{\cdot e}^{\left( -0.521\cdot T \right)}+4036$

Combining these two functions as per equation [1], we found that the relationship between constant temperature and the cost of development (*C*) was best described as

[8]	$C\left( T \right)=0.453\cdot T^{2}- 22.514\cdot T+293.927$

Whereby, *C* is minimised in *P. muralis* (14.5 ml CO_2_; Figure 1) when temperature (i.e., *T_opt_*) is 23.9 °C.

**Table S1. Sample sizes for *P. muralis* embryos incubated under one of three incubation treatments**. No variance (NV: 24 °C), low variance (LV: 22-26 °C), or high variance (HV: 18-30 °C) temperature treatments across different source populations (Region), and clutches (clutch order, 1^st^ or 2^nd^). Number of mothers for which response data was collected shown in parentheses.

| Response | Region | Incubation treatment | Sample size |
| --- | --- | --- | --- |
| 1. Metabolic rate (*MR*; ml CO_2_ h^-1^) |  |  |  |
|  | High | NV  LV  HV | 15 (10)  16 (10)  15 (9) |
|  | Mid | NV  LV  HV | 58 (27)  57 (27)  54 (26) |
|  | Low | NV  LV  HV | 24 (12)  27 (11)  28 (12) |
| 2. Development time (*D*; days) |  |  |  |
|  | High | NV  LV  HV | 1^st^ = 4 (4), 2^nd^ = 11 (10)  1^st^ = 7 (6), 2^nd^ = 12 (10)  1^st^ = 8 (6), 2^nd^ = 9 (9) |
|  | Mid | NV  LV  HV | 1^st^ = 13 (11), 2^nd^ = 33 (27)  1^st^ = 19 (13), 2^nd^ = 34 (27)  1^st^ = 19 (13), 2^nd^ = 32 (26) |
|  | Low | NV  LV  HV | 1^st^ = 7 (6), 2^nd^ = 14 (12)  1^st^ = 11 (8), 2^nd^ = 13 (11)  1^st^ = 11 (9), 2^nd^ = 14 (12) |
| 3. Cost of development (*C*; ml CO_2_) |  |  |  |
|  | High | NV  LV  HV | 15 (10)  16 (10)  15 (9) |
|  | Mid | NV  LV  HV | 58 (27)  57 (27)  54 (26) |
|  | Low | NV  LV  HV | 24 (12)  27 (11)  28 (12) |
| 4. Wet mass at hatching (g) |  |  |  |
|  | High | NV  LV  HV | 1^st^ = 4 (4), 2^nd^ = 11 (10)  1^st^ = 7 (6), 2^nd^ = 12 (10)  1^st^ = 8 (6), 2^nd^ = 9 (9) |
|  | Mid | NV  LV  HV | 1^st^ = 13 (11), 2^nd^ = 33 (27)  1^st^ = 19 (13), 2^nd^ = 34 (27)  1^st^ = 19 (13), 2^nd^ = 32 (26) |
|  | Low | NV  LV  HV | 1^st^ = 7 (6), 2^nd^ = 14 (12)  1^st^ = 11 (8), 2^nd^ = 13 (11)  1^st^ = 11 (9), 2^nd^ = 14 (12) |
| 5. Dry hatchling tissue mass (g) |  |  |  |
|  | High | NV  LV  HV | 15 (10)  16 (10)  15 (9) |
|  | Mid | NV  LV  HV | 58 (27)  57 (27)  54 (26) |
|  | Low | NV  LV  HV | 24 (12)  27 (11)  28 (12) |
| 6. Hatchling residual dry yolk mass (g) |  |  |  |
|  | High | NV  LV  HV | 15 (10)  16 (10)  15 (9) |
|  | Mid | NV  LV  HV | 58 (27)  57 (27)  54 (26) |
|  | Low | NV  LV  HV | 24 (12)  27 (11)  28 (12) |

**Table S2. Model ranking and selection**. Conditional Akaike Information Criterion (AICc) values were used to rank models for responses: 1. Metabolic rate (log_n_ ml CO_2_ h^-1^; for each temperature within incubation treatment: a. No variance (NV: 24 °C), b. Low variance (LV: 22 °C, 24 °C, 26 °C), c. High variance (HV: 18 °C, 24 °C, 30 °C), 2. Development time (log_n_ hours), 3. Cost of development (log_n_ ml CO_2_), 4. Hatchling wet mass (g), 5. Hatchling tissue dry mass (g), and 6. Hatchling residual dry yolk mass (g). Models for each response variable were ranked according to AICc value and ΔAICc is relative to best-fitting model (3 highest ranked models shown). Model parameters are provided in Table 1.

| Response | Model number | df | LogLik | AICc | ΔAICc | Weight |
| --- | --- | --- | --- | --- | --- | --- |
| 1a. Metabolic rate (NV) | 2 | 4 | 31.84 | -55.1 | 0.00 | 0.946 |
|  | 1 | 3 | 27.80 | -49.2 | 5.82 | 0.051 |
|  | 3 | 6 | 28.07 | -42.8 | 12.27 | 0.002 |
| 1b. Metabolic rate (LV) | 2 | 6 | 97.68 | -182.9 | 0.00 | 0.498 |
|  | 1 | 5 | 96.55 | -182.8 | 0.13 | 0.467 |
|  | 3 | 8 | 96.68 | -176.6 | 6.30 | 0.021 |
| 1c. Metabolic rate (HV) | 2 | 5 | 95.27 | -180.3 | 0.00 | 0.836 |
|  | 1 | 6 | 94.70 | -177.0 | 3.26 | 0.163 |
|  | 3 | 8 | 91.20 | -165.7 | 14.55 | 0.001 |
| 2. Development time | 7 | 5 | 636.46 | -1262.7 | 0.00 | 0.774 |
|  | 9 | 8 | 638.11 | -1259.8 | 2.95 | 0.177 |
|  | 8 | 6 | 634.26 | -1256.3 | 6.48 | 0.030 |
| 3. Cost of development | 8 | 6 | 97.01 | -181.6 | 0.00 | 0.869 |
|  | 7 | 5 | 94.05 | -177.8 | 3.80 | 0.130 |
|  | 9 | 8 | 92.57 | -168.4 | 13.19 | 0.001 |
| 4. Hatchling wet mass | 8 | 6 | 703.32 | -1394.4 | 0.00 | 1.000 |
|  | 9 | 8 | 695.43 | -1374.4 | 19.96 | 0.000 |
|  | 10 | 8 | 693.07 | -1367.6 | 26.79 | 0.000 |
| 5. Hatchling tissue dry mass | 8 | 6 | 788.18 | -1564.0 | 0.00 | 1.000 |
|  | 9 | 8 | 778.04 | -1539.4 | 24.56 | 0.000 |
|  | 16 | 10 | 772.34 | -1523.6 | 40.33 | 0.000 |
| 6. Hatchling yolk dry mass | 7 | 5 | 1212.12 | -2414.0 | 0.00 | 0.999 |
|  | 8 | 6 | 1206.35 | -2400.3 | 13.65 | 0.001 |
|  | 9 | 8 | 1191.89 | -2367.1 | 46.87 | 0.000 |

**Table S3. Summary of significance tests and marginal means (back transformed from log_n_) of fixed effects from linear mixed effect models.** Models describe the relationship between metabolic rate (log_n_ ml CO_2_ ml^-1^) and initial egg mass (log_n_ egg mass) at 24 °C (NV treatment only) across three stages through completion of embryonic development (days post oviposition; dpo): 25% (11-14 dpo), 50% (23-26 dpo), and 75% (36-39 dpo). Significant estimates shown in bold: **p* < 0.05, ***p* < 0.01, ****p* < 0.001. Letters represent significant pairwise differences across levels (a > b > c).

|  | χ^2^ | df | *p* | emmean (±SE) |
| --- | --- | --- | --- | --- |
| Intercept | 191.23 | 1 | **<0.001***** |  |
| Log_n_ Egg mass | 0.06 | 1 | 0.81 |  |
| Development stage | 6.54 | 2 | **0.04*** | 25%: 7.75⋅10^-3^ (2.83⋅10^-4^) ^c^  50%: 9.47⋅10^-2^ (2.79⋅10^-4^) ^b^  75%: 1.36⋅10^-2^ (4.10⋅10^-4^) ^a^ |
| Log_n_ Egg mass x Development stage | 1.12 | 2 | 0.57 |  |

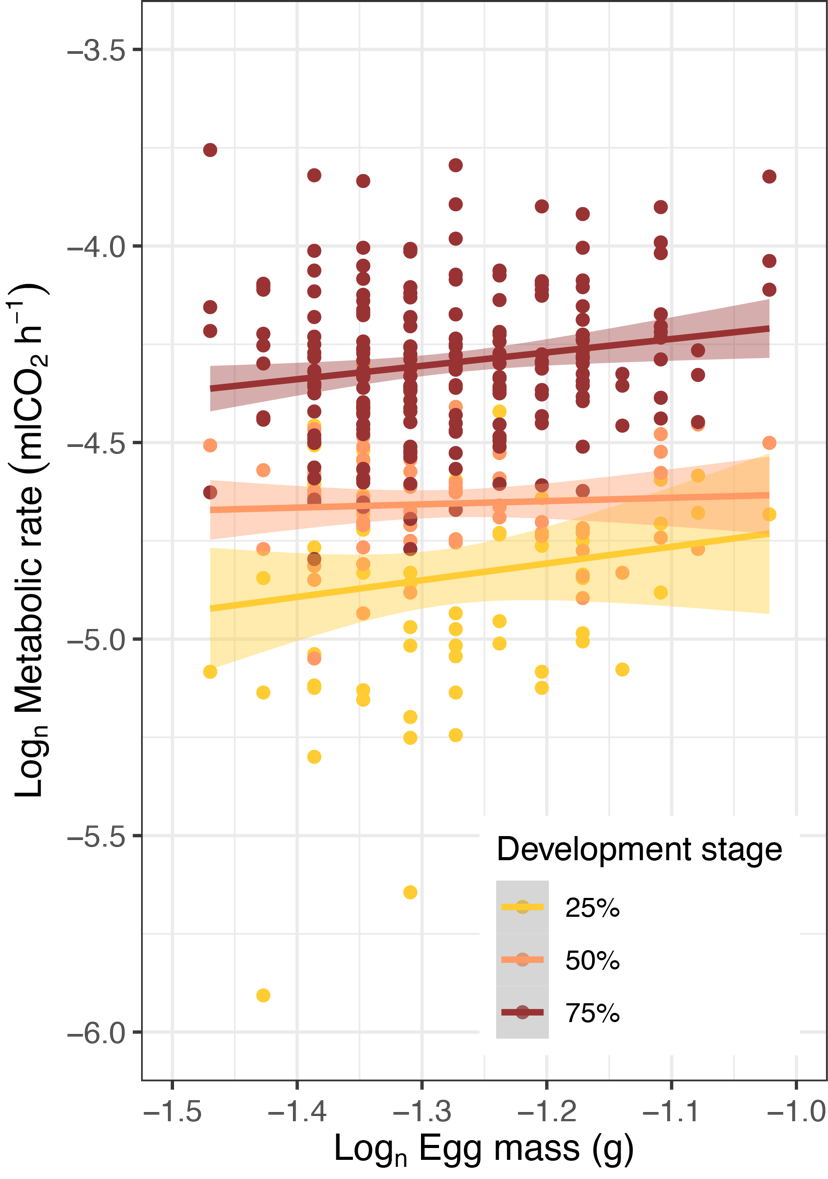

**Figure S1.** Relationship between Log_n_ egg mass and Log_n_ metabolic rate across embryonic development stages at T_opt_ (24 ºC) in *P. muralis* (25%, 50%, and 75% through completion from oviposition until hatching).
